# Small cytosolic dsDNAs repress cGAS activation and induce autophagy

**DOI:** 10.1101/2022.06.30.498186

**Authors:** Yao Liu, Xiao Chen, Yuemei Zhao, Xingyue Wang, Lina Chen, Weiyun Wang, Shouhui Zhong, Meizhen Hu, Zhizheng Dai, Jiayu Jiang, Xiaoxiao Cheng, Jiwei Zuo, Hui Liu, Di Ma, Xin Wang, Hongyu Ji, Shanshan Hu, Yu-Wei Luo, Ai-Long Huang, Kai-Fu Tang

## Abstract

Cyclic GMP-AMP (cGAMP) synthase (cGAS), a major cytosolic DNA sensor, activates innate immune responses by producing cGAMP, which activates stimulator of interferon genes (STING)^1^. Cytosolic DNA induces autophagy in a cGAS-dependent manner to avoid persistent immune stimulation. Although dsDNAs < 20 bp can bind to cGAS, robust cGAS activation requires dsDNAs > 45 bp ^2-4^. However, whether cytosolic dsDNAs < 45 bp exist in mammalian cells remains unclear. Here, we identified a class of small cytosolic DNAs (scDNAs) of ∼20–40 bp in human and mouse cell lines. scDNAs competed with herring testis DNA (HT-DNA, ∼200–1500 bp) for binding to cGAS, and repressing HT-DNA-induced cGAS activation and the associated interferon β (IFNβ) production. Moreover, scDNAs promoted cGAS and Beclin-1 interaction, triggering the release of Rubicon, a negative regulator of phosphatidylinositol 3-kinase class III (PI3KC3)^5,6^, from the Beclin-1–PI3KC3 complex, activating PI3KC3 and inducing autophagy. DNA damage decreased and autophagy inducers increased scDNA levels. scDNA transfection or autophagy induction attenuated DNA damage-induced cGAS-STING activation and IFNβ expression. Thus, scDNAs acted as molecular brakes of cGAS activation, preventing excessive inflammatory cytokine production following DNA damage. Our findings lay foundations for understanding the physiological and pathological functions of scDNAs.

---

When released into the cytoplasm, DNA fragments act as a danger signal triggering the innate immune system^1^. The cGAS-STING pathway has emerged as a major cell signaling pathway mediating the response to cytosolic dsDNAs. Binding of dsDNAs activates the catalytic activity of cGAS, leading to cGAMP synthesis from guanine and adenosine triphosphates (GTP and ATP). cGAMP binds to and activates STING, which in turn activates IκB kinase and TANK-binding kinase 1 and induces the expression of interferons and other cytokines^1^. Moreover, cytosolic dsDNAs induce autophagy to prevent excessive cGAS activation and persistent immune stimulation via two distinct mechanisms. First, dsDNAs induce autophagy by activating PI3KC3. Specifically, dsDNAs promote the interaction between cGAS and Beclin-1 and trigger the dissociation of Rubicon, a negative regulator of PI3KC3, from the Beclin-1–PI3KC3 complex^5,6,7^. Second, dsDNAs trigger cGAS-mediated cGAMP synthesis. cGAMP induces autophagy through a pathway dependent on STING but independent of the Beclin-1–PI3KC3 complex^8^. The immunostimulatory activity of cytosolic DNA depends on its length^1,9-11^. Long dsDNAs are more potent stimulators of cGAS than short dsDNAs^10^. *In vitro* studies revealed that although dsDNAs < 20 bp can bind to cGAS, robust activation of cGAS enzymatic activity requires longer dsDNAs > 45 bp^2-4^. Nevertheless, the following questions need to be answered: 1) Are there any dsDNAs < 45 bp in the cytoplasm of mammalian cells? 2) Could dsDNAs < 45 bp compete with longer dsDNA to bind cGAS and block cGAS activation? 3) Can dsDNAs < 45 bp induce autophagy in mammalian cells?

## Identification of small cytosolic dsDNAs

To determine whether dsDNAs < 45 bp exist in the cytoplasm, the cytosolic DNA purified from different human and mouse cell lines was subjected to electrophoresis in denaturing urea polyacrylamide gels. Interestingly, we found a class of small DNAs with apparent molecular sizes of ∼15–35 nt in the cytoplasm (Fig. 1a). S1 nuclease digestion slightly reduced the size of these small DNAs (Fig. 1b), whereas end-blunting with Klenow slightly increased their size (Fig. 1c), suggesting that these small DNAs were double-stranded DNAs with 5**′** protruding ends. After end-blunting, the scDNAs were cloned into the pGEM®-T Easy vector and subjected to deep sequencing. We found that the size distribution of the end-blunted scDNAs ranged from 20 to 40 bp (Fig. 1d and Extended Data Fig. 1a). Genome mapping showed that the scDNAs were randomly distributed on different chromosomes (Fig. 1e and Extended Data Fig. 1b). To investigate whether the scDNAs were nonspecific degradation products of cytosolic DNA during purification process, a ∼200 bp DNA fragment with no homology to human and mouse genomes was added to the cell lysis buffer immediately before cell lysis. We found that sequencing reads mapped to the spike-in DNA were undetectable, excluding the possibility that cytosolic dsDNAs were nonspecifically degraded into 20– 40 bp DNA fragments during DNA purification.

**Figure 1.**
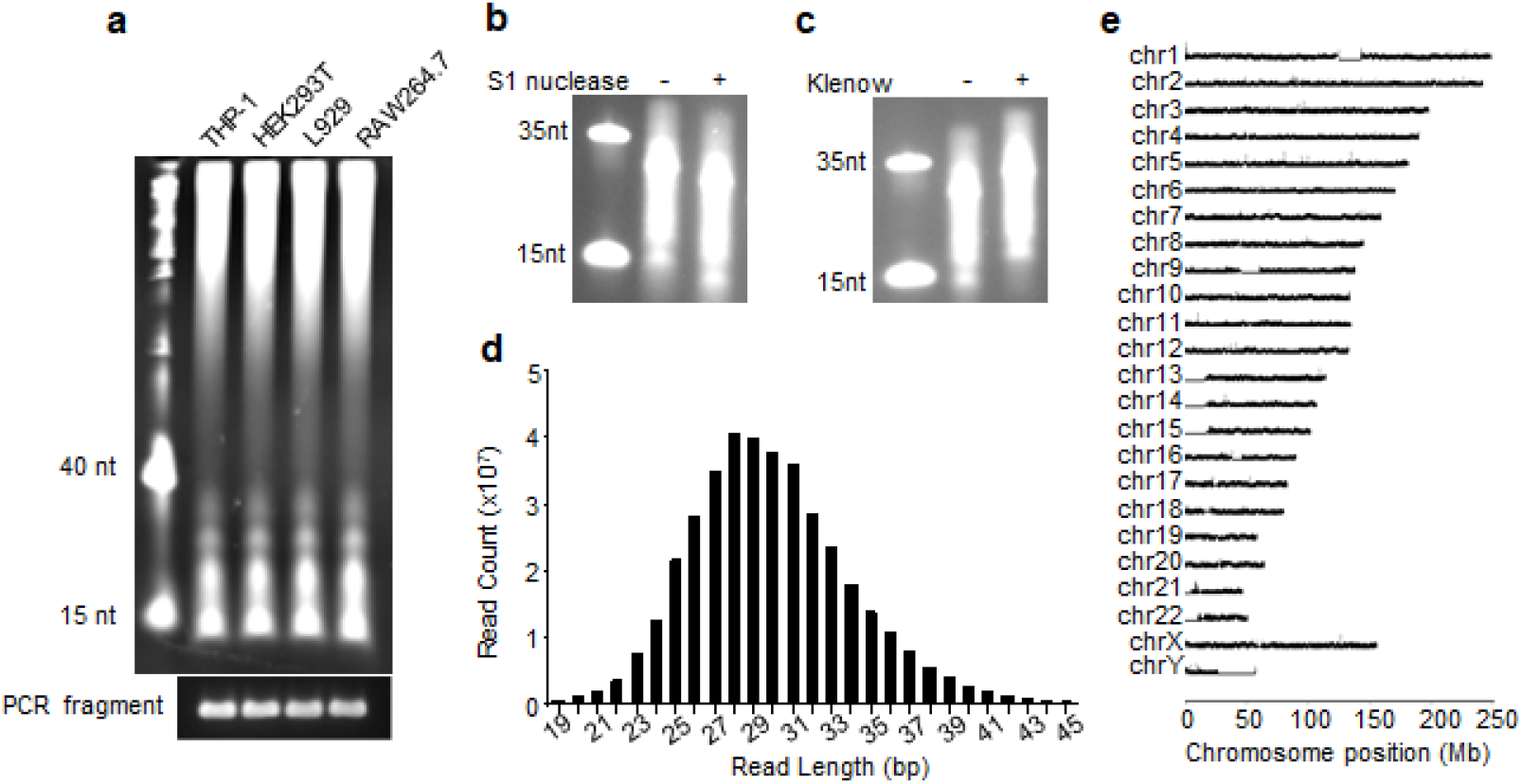
Identification and sequencing of small cytosolic DNAs (scDNAs). **a**, Polyacrylamide gel electrophoresis (PAGE) of cytosolic DNA purified from different cell types. The Polymerase chain reaction (PCR) products of spike-in DNA were used as loading controls. **b**, Purified scDNAs were digested with S1 nuclease and subjected to PAGE. **c**, Purified scDNAs were end-blunted with Klenow in the presence of dNTP and then subjected to PAGE. **d**, Read length distribution of scDNAs purified from THP-1 cells. Only the sequences that could be mapped to the genome are shown. **e**, Density of sequenced reads along different chromosomes is plotted as log_2_ (10 kilobases per million mapped reads).

## scDNAs blocked cGAS activation

We then investigated whether scDNAs could bind to and activate cGAS. *In vitro* biolayer interferometry assays demonstrated that scDNAs could bind to the recombinant cGAS protein (Fig. 2a). Transfection of RAW264.7 cells stably expressing enhanced green fluorescent protein (EGFP)-tagged cGAS with fluorescein-labeled scDNAs revealed that scDNAs formed puncta with cGAS in the cytoplasm (Fig. 2b). Although a 24 bp DNA duplex (P24) slightly induced interferon-β (IFNβ) expression and interferon regulatory factor 3 (IRF3) and STING phosphorylation, stimulation with purified scDNAs had no effect on IFNβ expression and IRF3 and STING phosphorylation. As a positive control, stimulation with herring testis DNA (HT-DNA) led to a considerable upregulation of IFNβ and an obvious phosphorylation of IRF3 and STING (Fig. 2c, d and Extended Data Fig. 2a, b). *In vitro* cGAS enzyme activity assays further confirmed that scDNAs could not robustly activate cGAS (Fig. 2e, f). These findings indicated that although scDNAs could bind to cGAS, they did not activate cGAS. We then investigated whether scDNAs could compete with long dsDNAs for binding to cGAS and inhibit cGAS activation upon stimulation with long dsDNAs. *In vitro* biolayer interferometry assays demonstrated that scDNAs could compete with HT-DNA to bind cGAS (Fig. 2g and Extended Data Fig. 2c). Co-transfection experiments showed that scDNAs repressed HT-DNA-induced IFNβ expression and IRF3 and STING phosphorylation (Fig. 2h, i and Extended Data Fig. 2d–g). *In vitro* cGAS enzyme activity assays confirmed that scDNAs repressed HT-DNA-induced cGAS activation (Fig. 2e, f). Overall, these data indicated that scDNAs competed with long dsDNAs and reduced cGAS activation. Further experiments demonstrated that the effect of scDNAs on cGAS activation was not cell- or species-specific, as scDNAs purified from mouse cells could inhibit HT-DNA-induced IFNβ expression and IRF3 and STING phosphorylation in human cells and vice versa (Extended Data Fig. 2d–g).

**Figure 2.**
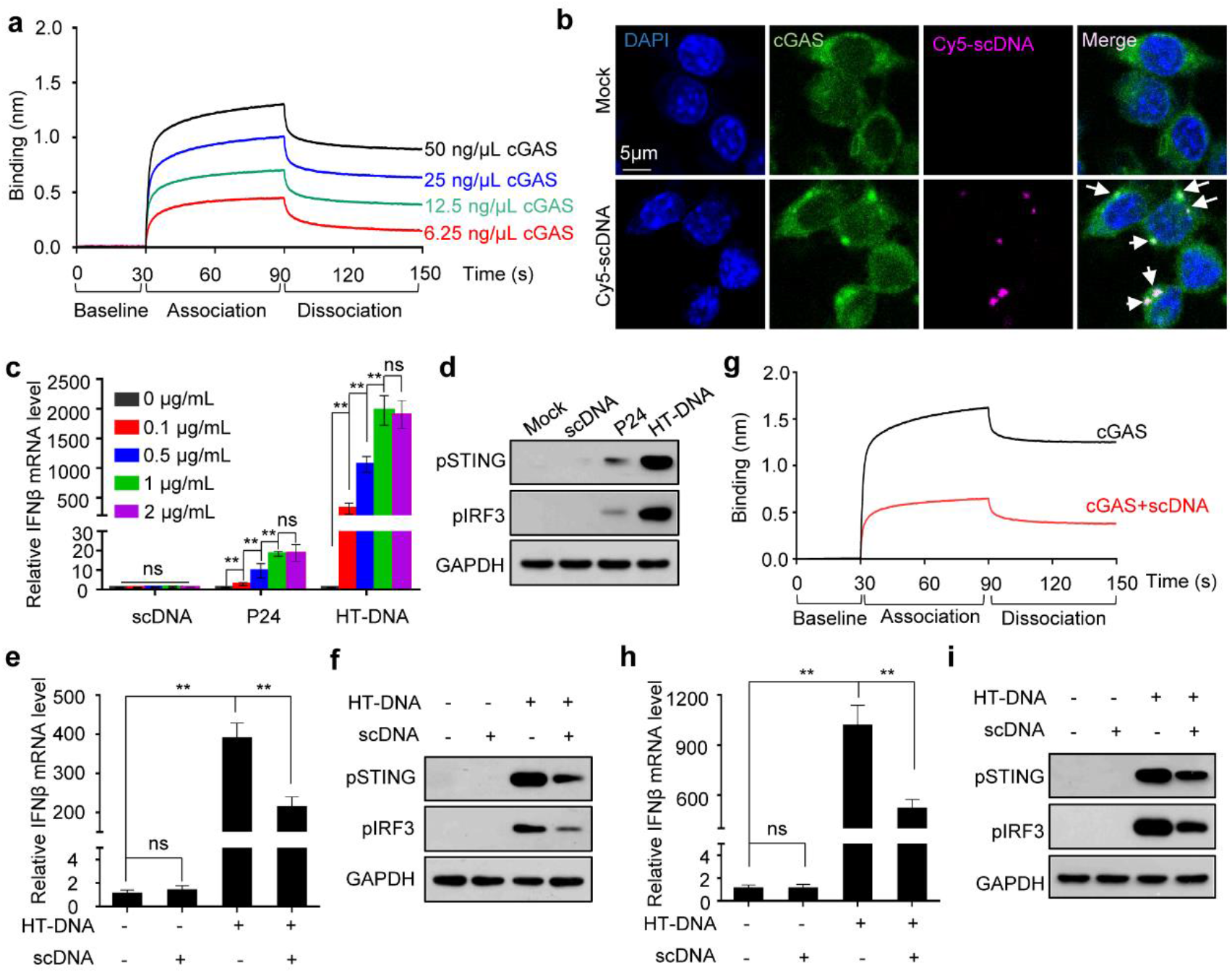
scDNAs compete with long dsDNA to bind cyclic GMP-AMP synthase (cGAS) and inhibit long dsDNA-induced cGAS activation. **a**, Representative scDNAs-cGAS biolayer interferometry sensorgram. Different concentrations of recombinant cGAS protein were tested for binding to biotin-labeled scDNAs immobilized on a streptavidin-coated biosensor. **b**, RAW264.7 cells stably expressing EGFP-cGAS were transfected with Cy5-labeled scDNAs and analyzed using confocal microscopy. Arrows indicate the colocalization of scDNAs and cGAS puncta. **c**, RAW264.7 cells were transfected with scDNAs, a 24 bp dsDNA (P24), or herring testis DNA (HT-DNA) at different concentrations, and interferon β (*IFNβ*) mRNA levels were quantified. **d**, RAW264.7 cells were transfected with scDNAs, P24, or HT-DNA (1 µg/mL) or mock-transfected, and the levels of phosphorylated (p) stimulator of interferon genes (STING) and p-interferon regulatory factor 3 (IRF3) were determined. **e, f**, Recombinant cGAS protein was incubated with ATP, GTP, HT-DNA, scDNAs, or both HT-DNA and scDNAs. The relative cGAMP amount in the supernatant was determined after its delivery into perfringolysin O-permeabilized L929 cells, followed by measurement of *IFNβ* mRNA (e), pSTING, and pIRF3 (f) levels. cGAS incubated with ATP and GTP, but without DNA, was used as control. **g**, Representative biolayer interferometry sensorgram showing the competition of scDNAs and HT-DNA for binding to cGAS. Biotin-labeled HT-DNA was immobilized on a streptavidin-coated sensor and then loaded with cGAS alone (50 ng/µL) or cGAS pre-incubated with unlabeled scDNAs (300 ng/µL). **h, i**, RAW264.7 cells were transfected with HT-DNA (1 µg/mL) or scDNAs (1 µg/mL) or co -transfected with both HT-DNA (1 µg/mL) and scDNAs (1 µg/mL), and *IFNβ* mRNA (h), pSTING, and pIRF3 (i) levels were determined. Data (c, e, h) are expressed as mean ± standard error of the mean (s.e.m) of three biological replicates. Statistical analysis was performed using two-sided Student’s *t*-tests; ** indicates p < 0.01; ns indicates p > 0.05.

## scDNA-induced autophagy

We observed that scDNAs had no effect on cGAS activity, suggesting that scDNAs could not activate autophagy by inducing cGAMP production. We, therefore, explored whether scDNAs could induce autophagy by releasing Rubicon from the Beclin-1– PI3KC3 complex. Co-immunoprecipitation experiments revealed that similar to HT-DNA stimulation, scDNAs enhanced the interaction between cGAS and the Beclin-1– PI3KC3 complex and decreased the interaction between Rubicon and the Beclin-1– PI3KC3 complex (Fig. 3a). As Rubicon inhibits PI3KC3 kinase activity^5,6^, we investigated the effect of scDNAs on PI3KC3 enzymatic activity by measuring intracellular phosphatidylinositol-3-phosphate (PtdIns-3-P) levels and distribution. We used the p40PX-EGFP fusion protein as a non-invasive probe because the PX domain of p40phox specifically binds PtdIns-3-P and forms a punctate cytoplasmic pattern in the presence of PtdIns-3-P-rich vesicles^12^. Stimulation with scDNAs led to a considerable increase in the number of p40PX-EGFP puncta (Fig. 3b). The number of scDNA-induced p40PX-EGFP puncta considerably decreased following cGAS depletion in RAW264.7 cells and strongly increased upon cGAS expression in HEK293T cells (Extended Data Fig. 3a, b). To further confirm that the scDNA-induced increase in PI3KC3 enzymatic activity was dependent on cGAS expression, we overexpressed Beclin-1 and PI3KC3 (Vps34) in the presence or absence of cGAS in HEK293T cells and measured PI3KC3 activity after scDNAs stimulation. Consistent with a previous report^7^, ectopic cGAS expression enhanced PI3KC3 activity, and HT-DNA stimulation further augmented PtdIns-3-P production (Extended Data Fig. 3c). Moreover, scDNAs stimulated PtdIns-3-P production in cGAS-expressing cells, but not in cells that do not express cGAS (Extended Data Fig. 3c). Because PtdIns-3-P is essential for autophagy initiation^7,13^, we investigated whether scDNAs could induce autophagy. Our results revealed that scDNAs stimulated the conversion of LC3 into a lipidated form (LC3-II) and the formation of EGFP-LC3 punctate structures, which were more prominent when cells were pre-treated with chloroquine (Fig. 3 c, d and Extended Data Fig. 4 a–d). Furthermore, electron microscopy demonstrated that scDNAs induced autophagic vacuole formation (Fig. 3e). Although scDNA treatment increased *p62* mRNA levels (Extended Data Fig. 4e), p62 protein levels were reduced (Fig. 3c and Extended Data Fig. 4a, b). These data indicated that scDNAs activated autophagy. scDNA-induced LC3 lipidation and formation of EGFP-LC3 puncta were obviously reduced following cGAS depletion in RAW264.7 cells (Extended Data Fig. 5a, b) and increased upon cGAS expression in HEK293T cells (Extended Data Fig. 5c, d). These data indicated that cGAS was required for scDNA-induced autophagy. The effect of scDNAs on autophagy was not cell- or species-specific, as scDNAs purified from human cells could induce LC3 lipidation in mouse cells and vice versa (Extended Data Fig. 4a, b). These findings indicated that similar to long cytosolic DNA, scDNAs could induce autophagy by promoting the interaction between cGAS and Beclin-1.

**Figure 3.**
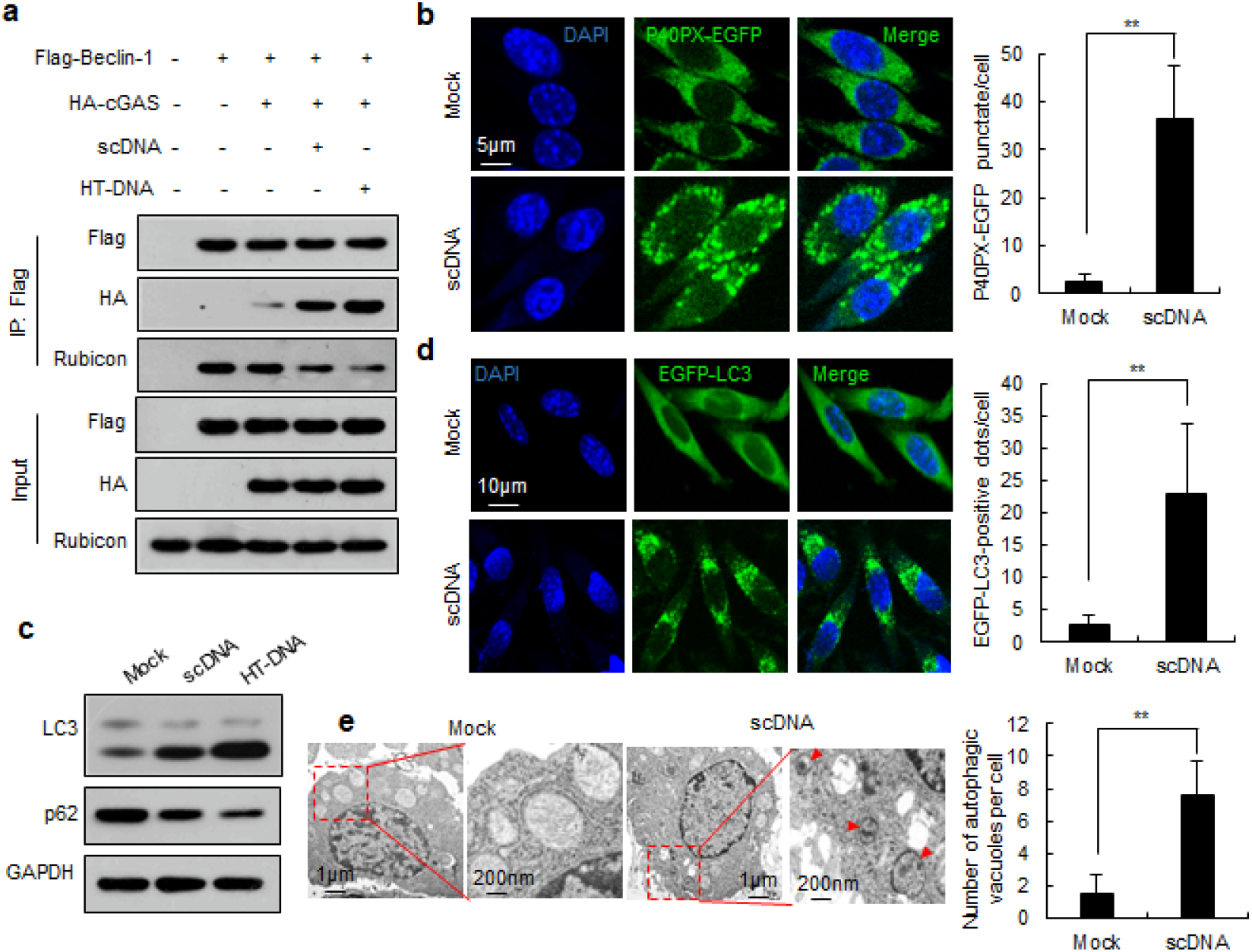
scDNAs induce autophagy by releasing Rubicon from the Beclin-1-phosphatidylinositol 3-kinase class III (PI3KC3) complex. **a**, HEK293T-cGAS or HEK293T-Vector cells were transfected with Flag-Beclin-1 plasmid and stimulated with scDNAs or HT-DNA (1 µg/mL). The cell lysates were subjected to immunoprecipitation and immunoblotting with the indicated antibodies. **b**, RAW264.7 cells stably expressing p40PX-EGFP were stimulated with scDNAs (1 µg/mL) or mock-transfected. The number of p40PX-EGFP foci was determined using confocal microscopy. Data are expressed as mean ± s.e.m of three biological replicates, *n* ≥ 200 cells. Statistical analysis was performed using two-sided Student’s *t*-tests; ** indicates p < 0.01. **c**, L929 cells were transfected with scDNAs (1 µg/mL) or mock-transfected, and LC3-II and p62 levels were determined. HT-DNA was used as a positive control. **d**, L929 cells stably expressing EGFP-LC3 were transfected with scDNAs (1 µg/mL) or mock-transfected and analyzed using confocal microscopy. Data are expressed as mean ± s.e.m of five biological replicates, *n* ≥ 200 cells. Statistical analysis was performed using two-sided Student’s *t*-tests; ** indicates p < 0.01. **e**, Representative electron micrographs of autophagic vacuoles (indicated by arrows) in RAW264.7 cells stimulated with 1 µg/mL scDNAs and the number of autophagic vacuoles per cell in cells stimulated with (n = 52) or without scDNAs (n = 50). Data are expressed as mean ± s.e.m of three biological replicates. Statistical analysis was performed using two-sided Student’s *t*-tests; ** indicates p < 0.01.

## DNA damage decreased scDNA levels

The nucleotide excision repair (NER) pathway removes ultraviolet (UV) photoproducts from the cellular genome in the form of ∼20–30 nt oligonucleotides^14^. If these small DNAs were byproducts of NER, we expected that their levels would increase upon UV irradiation. However, UV irradiation decreased the levels of these small DNAs (Fig. 4a and Extended Data Fig. 6a), suggesting that they were unlikely to be byproducts of NER. Having shown that UV radiation decreased scDNA levels, we investigated whether other DNA-damaging agents could affect scDNA levels. Treatment with etoposide, adriamycin, and camptothecin also reduced scDNA levels (Fig. 4b and Extended Data Fig. 6b, c). DNA damage induces the presence of cytosolic DNA, which activates the cGAS-STING pathway and induces IFNβ expression^15^. Consistently, we found that exposure to UV radiation, camptothecin, etoposide, and adriamycin led to cytosolic dsDNA accumulation, STING and IRF3 phosphorylation, and IFNβ upregulation (Extended Data Fig. 6d–i). Transfection with scDNAs partially blocked STING and IRF3 phosphorylation and IFNβ upregulation induced by DNA-damaging agents (Fig. 4c, d and Extended Data Fig. 7). These data indicated that both the increase in long cytosolic dsDNAs and the decrease in scDNAs contributed to DNA damage-induced activation of the cGAS-STING-IRF3 pathway.

**Figure 4.**
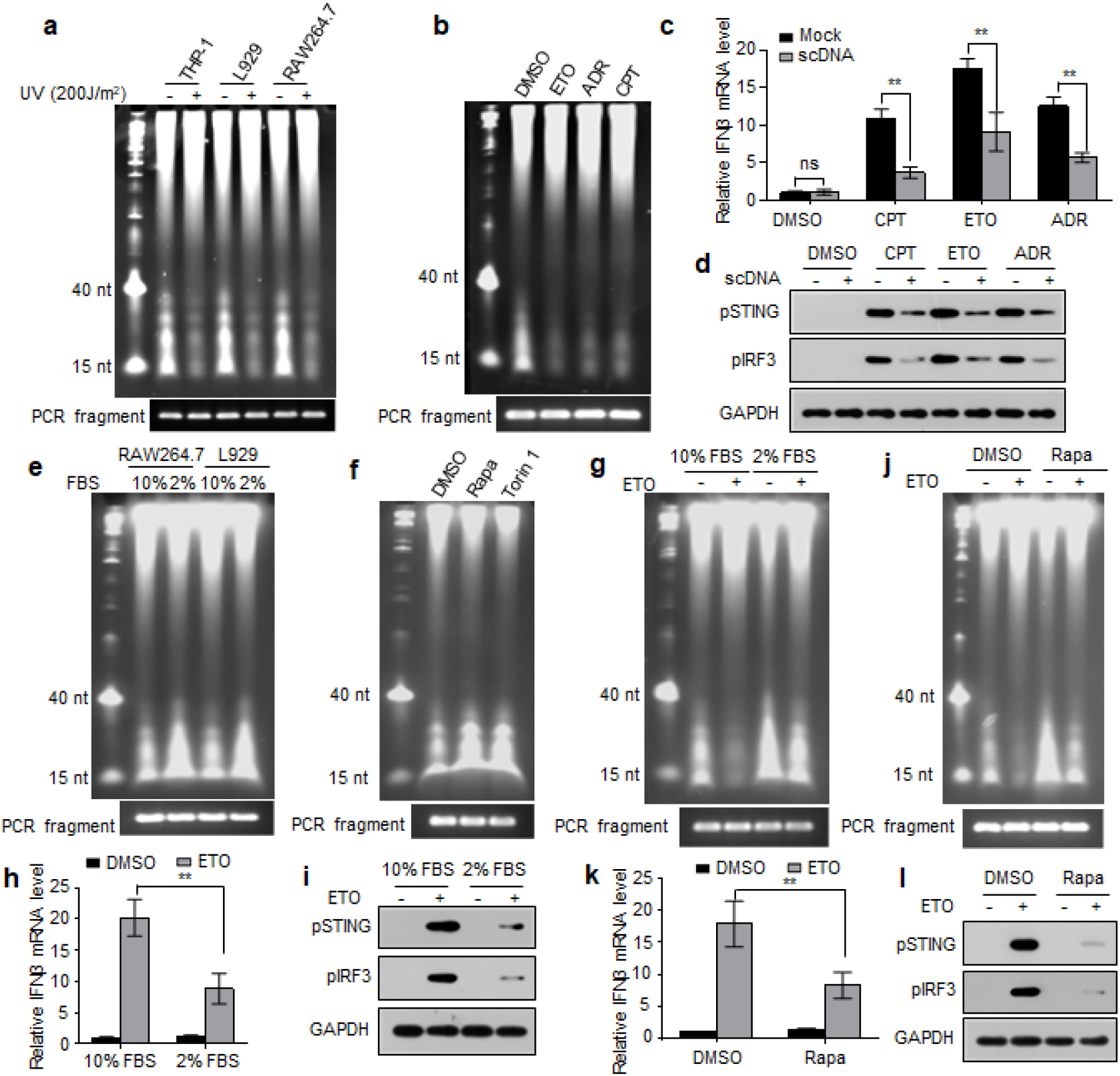
Downregulation of scDNAs facilitates cGAS-STING activation following DNA damage. **a**, Different cells were irradiated with 200 J/m^2^ ultraviolet (UV) light or unirradiated, and cytosolic DNA was subjected to PAGE analysis. **b**, RAW264.7 cells were treated with etoposide (ETO, 2 µM), adriamycin (ADR, 5 µM), or camptothecin (CPT, 0.2 µM), and cytosolic DNA was subjected to PAGE analysis. **c, d**, RAW264.7 cells were transfected with scDNAs (0.25 µg/mL) or mock-transfected and exposed to camptothecin (0.2 µM), etoposide (2 µM), or adriamycin (5 µM), and *IFNβ* mRNA (c), pSTING, and pIRF3 (d) levels were determined. **e**, RAW264.7 and L929 cells were exposed to 10% or 2% fetal bovine serum (FBS), and cytosolic DNA was subjected to PAGE analysis. **f**, RAW264.7 cells were exposed to rapamycin (Rapa, 2 µM) or torin1 (0.1 µM), and cytosolic DNA was subjected to PAGE analysis. **g**, RAW264.7 cells were exposed to 10% or 2% FBS in the presence or absence of etoposide (2 µM), and cytosolic DNA was subjected to PAGE analysis. **h, i**, RAW264.7 cells were exposed to 10% or 2% FBS in the presence or absence of etoposide (2 µM), and *IFNβ* mRNA (h), pSTING, and pIRF3 (i) levels were determined. **j**, RAW264.7 cells were treated with etoposide (2 µM) in the presence or absence of rapamycin (2 µM), and cytosolic DNA was subjected to PAGE analysis. **k, l**, RAW264.7 cells were treated with etoposide (2 µM) in the presence or absence of rapamycin (2 µM), and *IFNβ* mRNA (k), pSTING, and pIRF3 (l) levels were determined. The PCR products of spike-in DNA were used as loading controls (a, b, e–g, j). Data (c, h, k) are expressed as mean ± s.e.m of three biological replicates. Statistical analysis was performed using two-sided Student’s *t*-tests; ** indicates p < 0.01; ns indicates p > 0.05.

## Autophagy inducers upregulated scDNAs

DNA damage can induce autophagy^16^, which inhibits the cGAS-STING-IRF3 signaling pathway by degrading cGAS, STING, and IRF3 and inhibiting their activities, or by degrading cytosolic DNA^7,17-19^. However, whether scDNAs are involved in the regulation of cGAS-STING-IRF3 signaling by autophagy remains unclear. We found that serum deprivation or treatment with rapamycin or torin1 (two mTOR inhibitors widely used to induce autophagy) increased scDNA levels (Fig. 4e, f and Extended Data Fig. 8a, b). Moreover, serum deprivation or rapamycin treatment not only prevented the decrease in scDNA levels induced by DNA damage, but also alleviated DNA damage-induced STING and IRF3 phosphorylation and IFNβ upregulation (Fig. 4g–l and Extended Data Fig. 8c–h). These data suggested that autophagy inducers could restrict DNA damage-induced cGAS-STING activation by upregulating scDNAs.

## Discussion

Research interest in extrachromosomal DNA molecules, such as extrachromosomal circular DNA and cytosolic DNA, has been increasing during the last decade^1,20^. Here, we identified a new class of small cytosolic dsDNAs in human and mouse cell lines, which we propose to call scDNAs. Functional analysis revealed that scDNAs repressed long dsDNA-induced cGAS-STING activation and induced autophagy in a cGAS-dependent manner, indicating that scDNAs served as molecular brakes for cGAS activation that restricted inflammation induced by cytosolic DNA. Previous studies have revealed that DNA damage activates the cGAS-STING pathway by inducing the presence of long cytosolic dsDNAs^15^. We found that DNA-damaging treatment decreased scDNA levels. Transfection with purified scDNAs partially blocked DNA damage-induced cGAS-STING activation. These results indicated that DNA damage-induced scDNA downregulation facilitated cGAS activation. DNA damage induces autophagy, which represses DNA damage-induced cGAS-STING activation directly by inhibiting and degrading cGAS, STING, and IRF3 or indirectly by clearing cytosolic DNA^7,16-19^. In the present study, we found that autophagy inducers increased scDNA levels and repressed DNA damage-induced cGAS activation. Our findings implied that autophagy inducers could block DNA damage-induced cGAS activation by upregulating scDNAs; however, methods to specifically decrease scDNA levels are lacking, rendering it difficult to establish such a conclusion with confidence.

This study raises the following questions. What is the molecular mechanism underlying scDNA biogenesis? Why were scDNAs downregulated following DNA-damaging treatment? How do autophagy inducers upregulate scDNAs? Can other physiological and pathological stimuli regulate scDNA expression levels? Given that scDNAs not only existed in cGAS-expressing cells (RAW264.7, THP-1, and L929 cells) but also in cells that do not express cGAS (HEK293T cells), they might contribute to cGAS-independent cellular functions. Therefore, further studies should investigate whether scDNAs can bind to other DNA sensors (AIM2, DAI, IFI16, among others^21^) and modulate their functions. Furthermore, future studies should also evaluate the therapeutic implications of scDNAs in the treatment of cytosolic DNA-associated inflammatory diseases.

## Methods

### Cell culture and treatment

The human embryonic kidney cell line HEK293T, human monocyte cell line THP-1, mouse fibrosarcoma cell line L929, and mouse macrophage cell line RAW264.7 were purchased from the American Type Culture Collection (Manassas, VA, USA). HEK293T and RAW264.7 cells were grown in Dulbecco’s Modified Eagle Medium (Hyclone, Logan, UT, USA) supplemented with 10% fetal bovine serum (FBS). THP-1 cells were cultured in Roswell Park Memorial Institute (RPMI) 1640 medium supplemented with 10% FBS and 50 μM β-mercaptoethanol (Sigma-Aldrich, St. Louis, MO, USA). L929 cells were cultured in RPMI 1640 medium supplemented with 10% FBS. Cells were cultured at 37 °C in a 5% CO_2_ humidified incubator. All the cell lines were mycoplasma-free and authenticated by the Genetic Testing Corporation (Suzhou, China) using polymorphic short tandem repeat loci.

For UV irradiation, cells were washed with phosphate-buffered saline (PBS) and then exposed to the indicated doses of UV using a lamp emitting 254 nm light. After UV irradiation, cells were replenished with a culture medium and incubated at 37 °C for 4 h. To induce DNA damage, cells were exposed to different concentrations of camptothecin, etoposide, or adriamycin for 24 h. For autophagy induction, cells were cultured in media supplemented with 2% FBS, or exposed to rapamycin (MedChemExpress, Monmouth Junction, NJ, USA) or torin 1 (MedChemExpress) for 6 h. To investigate the combined effects of UV irradiation and autophagy induction, UV irradiated cells were cultured in media supplemented with 2% FBS or exposed to rapamycin for 6 h. To investigate the combined effects of etoposide (MedChemExpress) and autophagy induction, cells were cultured in media supplemented with etoposide and 2% FBS, or in media supplemented with etoposide, 10% FBS and rapamycin for 24 h.

### DNA and siRNA transfection

Cells were transfected with plasmids, small interfering RNAs (siRNAs), herring testis DNA (HT-DNA) (Sigma-Aldrich), a 24 bp dsDNA (P24, listed in Extended Data Tables 1), or small cytosolic DNAs (scDNAs) using Lipofectamine 2,000 (Thermo Fisher Scientific Inc., Waltham, MA, USA) according to the manufacturer’s instructions. The medium was changed to complete growth media 4 h after plasmid or siRNA transfection or 1 h after HT-DNA or scDNA transfection. To investigate the effects of P24, HT-DNA or scDNAs on cyclic GMP-AMP synthase (cGAS)-stimulator of interferon genes (STING) activation, cells were harvested 9 h after transfection. To investigate the effects of HT-DNA or scDNAs on phosphatidylinositol 3-kinase (PI3K) activation and autophagy induction, cells were harvested 6 h after transfection. To investigate the effect of scDNA transfection on UV-induced cGAS activation, cells were irradiated with UV 1 h after transfection and harvested 4 h after UV irradiation. To investigate the effect of scDNA transfection on etoposide-induced cGAS activation, cells were transfected with scDNAs at 18 h after etoposide treatment, and harvested 6 h after transfection. The pEGFP-N1-cGAS plasmid was constructed by inserting the coding sequence of the human *cGAS* into the pEGFP-N1 vector and was customized by Inovogen Tech. Co. (Beijing, China). The other plasmids and siRNAs used are listed in Extended Data Tables 2 and 3.

### Lentiviral preparation

For lentivirus preparation, HEK293T cells were transiently transfected with a lentiviral vector together with pMD2.G and psPAX2 using Lipofectamine 2,000. After 2–3 d of culture, the supernatant was passed through a 0.45 µm syringe filter unit (SLHV033RB; EMD Millipore, Burlington, MA, USA). HBLV-EGFP-LC3-PURO lentivirus was purchased from Hanbio Biotechnology (Shanghai, China).

### scDNA identification and sequencing

Cells were resuspended in cell lysis buffer (10 mM HEPES, pH 7.9, 10 mM KCl, 1.5 mM MgCl_2_, 0.5 mM β-mercaptoethanol) and incubated for 15 min on ice. Nonidet P-40 was then added to a final concentration of 0.2%, and the cells were incubated for 5 min on ice, followed by low-speed centrifugation at 1,750 × *g* at 4 °C for 5 min. The supernatant (i.e., the cytosolic fraction) was centrifuged at 14,000 × *g* at 4 °C for 15 min to remove cell debris and insoluble aggregates. RNAs in the cytosolic fraction were removed by treatment with 20 µg/mL RNase A (Sigma-Aldrich) at 37 °C for 6 h, and the proteins were then digested with 10 µg/mL proteinase K (Sigma-Aldrich) at 55 °C overnight. DNA was recovered by phenol/chloroform and chloroform extraction, followed by isopropanol precipitation. The purified cytosolic DNAs were subjected to 6% denaturing urea polyacrylamide gel electrophoresis (PAGE). To ensure equal loading, a fragment of the pGEM®-T Easy vector (Promega, Madison, WI, USA) was spiked into the cytosolic fraction according to the protein concentration. Before gel electrophoresis, DNA concentration was measured using a NanoDrop spectrophotometer (ND-1000, NanoDrop Technologies, Wilmington, DE, USA), and spike-in DNA was quantified by real-time polymerase chain reaction (PCR) using the Teasy primers listed in Extended Data Table 1. The small DNA fragments in the cytosolic fraction were purified using the PAGE gel DNA/RNA purification kit (BioTeke, Beijing, China) according to the manufacturer’s instructions. To characterize scDNAs, gel-purified scDNAs were digested with S1 nuclease (Thermo Fisher Scientific Inc.), or end-blunted with Klenow (New England Biolabs, Inc. Beverly, MA, USA) according to the manufacturer’s instructions. To sequence scDNAs, the gel-purified scDNAs were converted into repaired DNA with 5′-phosphorylated and 3′-dA-tailed ends using the End Prep enzyme mix from Vazyme Biotech Co., Ltd. (Nanjing, China), according to the manufacturer’s instructions. The repaired DNA was ligated with the pGEM®-T Easy vector and then PCR-amplified using the Teasy primers listed in Extended Data Table 1. The PCR product was used to construct libraries and sequenced on an Illumina NovaSeq 6,000 (Illumina, San Diego, CA, USA) with 150 bp paired-end reads. Low-quality sequencing reads were discarded, and sequences derived from the pGEM®-T Easy vector were removed using the Cutadapt program (Version 1.16)^22^. Then, the sequences were mapped to the human (GRCh38/hg38) and mouse (GRCm38/mm10) reference genomes using HISAT2 (Version 2.1.0)^23^ with default parameters and subsequently processed using samtools^24^. Normalized read counts in each 10 kb window on the chromosome were obtained using bedtools (Version 2.27.1)^25^.

### *In vitro* cGAS activity assay

Recombinant human cGAS was prepared as previously described^26^. The *in vitro* cGAS reaction was performed as previously described^4^ with modifications. Briefly, recombinant human cGAS protein was mixed with ATP (5 mM), GTP (300 μM), HT-DNA (15 ng/μL), or/and scDNAs (15 ng/μL) in a reaction buffer (20 mM Tris-hydrogen chloride [HCl] pH 7.5, 5 mM MgCl_2_ and 0.2 mg/mL bovine serum albumin). After incubation at 37 °C for 2 h, the reaction was terminated by heating at 95 °C for 5 min. After centrifugation at 20,000 × *g* for 5 min, the supernatant was transferred to L929 cells permeabilized with 50 ng/mL perfringolysin O (Cusabio, Wuhan, China). The cells were cultured at 37 °C for 9 h and then harvested for measurement of IFNβ expression.

### Biolayer interferometry assay

scDNAs binding to cGAS was assessed by biolayer interferometry^27^ using the BLItz system (ForteBio, Menlo Park, CA, USA). HT-DNA and scDNAs were biotin-labeled using EZ-Link Hydrazide Biotin (#21339, Thermo Fisher Scientific Inc.), EDC (#22980, Thermo Fisher Scientific Inc.), and imidazole according to the manufacturer’s instructions. The biotin-labeled scDNAs or HT-DNA (25 μg/mL) was immobilized onto a ForteBio streptavidin biosensor overnight at 4 °C. The biosensor was washed in binding buffer [20 mM Tris-HCl, pH 7.5, and 150 mM sodium chloride (NaCl)] for 30 s to remove unbound DNA, and purified cGAS diluted in binding buffer or a binding buffer vehicle control was loaded onto the biosensor for 60 s (i.e., the association step). To measure dissociation, the biosensor was washed with binding buffer for 60 s. To measure the competition between scDNAs and HT-DNA for cGAS binding, cGAS was premixed with unlabeled HT-DNA or scDNAs for 1 h at 4 °C, and the premix was then presented to the biosensor that had already been loaded with biotin-labeled scDNAs or HT-DNA.

### Western blotting

Cell lysates were prepared in radioimmunoprecipitation assay buffer supplemented with protease inhibitor cocktail tablets (Roche, Basel, Switzerland), and total protein was measured using a Bicinchoninic Acid Protein Assay Kit (Beyotime Biotechnology, Shanghai, China). Equal amounts of protein were resolved by sodium dodecyl sulfate-PAGE and transferred to a polyvinylidene fluoride membrane (Bio-Rad Laboratories, Hercules, CA, USA). Blots were probed with the indicated primary antibodies listed in Extended Data Table 4, followed by incubation with appropriate horseradish peroxidase-conjugated secondary antibodies and detection using Clarity Western enhanced chemiluminescence substrate (Bio-Rad Laboratories).

### Immunoprecipitation

HEK293T-cGAS cells (HEK293T stably expressing cGAS) were generated by infecting HEK293T cells with lentivirus expressing HA-tagged cGAS. HEK293T-Vector cells were generated by infecting HEK293T cells with control lentivirus. HEK293T-cGAS and HEK293T-Vector cells were maintained in the presence of puromycin (2 µg/mL, Hyclone). HEK293T-cGAS or HEK293T-Vector cells were transfected with the Flag-Beclin-1 plasmid. Forty-eight hours after plasmid transfection, cells were transfected with or without HT-DNA or scDNAs (1 μg/mL) and incubated at 37 °C for 6 h. Cells were then lysed with immunoprecipitation (IP) buffer (20 mM HEPES, pH 7.4, 150 mM NaCl, 2 mM MgCl_2_, 0.5% Triton X-100, 0.1% Tween-20) with a protease inhibitor cocktail at 4 °C for 30 min with continuous rotation and then centrifuged at 13,000 × *g* for 10 min. Equal amounts of lysate were immunoprecipitated with anti-Flag M2 affinity gel (Sigma-Aldrich) at 4 °C overnight. The gel was then washed three times with 1.5 mL Tris-buffered saline (50 mM Tris-HCl, 150 mM NaCl, pH 7.4) and eluted with 0.1 M glycine (pH 3.0) according to the manufacturer’s instructions. The eluates were immediately neutralized with 1 M Tris (pH 8.0) and subjected to western blot analysis.

### PI3KC3 kinase assay

The phosphatidylinositol 3-kinase class III (PI3KC3) (hVps34) kinase assay was performed as previously described^7^. Briefly, FLAG-tagged hVps34 and Myc-tagged Beclin-1 plasmids were co-transfected into HEK293T-cGAS and HEK293T-Vector cells; 48 h later, the cells were transfected with HT-DNA or scDNAs (1 µg/mL) or mock-transfected. Cells were harvested 6 h after the second transfection and lysed using IP buffer. Vps34 complexes were immunoprecipitated with anti-Flag M2 affinity gel (Sigma-Aldrich) and washed successively with IP buffer and 2X reaction buffer (20 mM Tris pH 8, 200 mM NaCl, 2 mM ethylenediamine tetraacetic acid, 20 mM manganese(II) chloride, and 100 μM ATP). The PI3KC3 kinase activity assay and the production of phosphatidylinositol (3,4)-bisphosphate [PI(3,4)P2] were measured using a Class III PI3-Kinase Kit according to the manufacturer’s protocols (Echelon Biosciences Inc. Salt Lake City, UT, USA).

### Confocal microscopy

#### cGAS–DNA puncta imaging

RAW264.7 EGFP-cGAS cells (RAW264.7 cells stably expressing EGFP-tagged cGAS) were established by transfection with pEGFP-N1-cGAS plasmid and selected with 1 mg/mL G418. scDNAs were Cy5-labeled using the Label IT® Nucleic Acid Labeling Kit (Mirus Bio LLC, Madison, WI, USA; WI 53719), according to the manufacturer’s instructions. To investigate the formation of cGAS-scDNAs foci within the cells, RAW264.7 EGFP-cGAS cells were seeded on coverslips in 24-well culture dishes and transfected with Cy5-labeled scDNAs. Two hours after transfection, cells were fixed with 4% paraformaldehyde in PBS and imaged using laser scanning confocal microscopy (LEICA DMi8, Leica Microsystems, Wetzlar, Germany).

#### LC3 puncta imaging

L929, RAW264.7, HEK293T-cGAS, and HEK293T-Vector cells were infected with HBLV-EGFP-LC3-PURO lentivirus, and transduced cells were selected with puromycin (L929 cells: 4 µg/mL; RAW264.7 cells: 1 µg/mL ; HEK293T-cGAS and HEK293T-Vector cells: 8 µg/mL). To investigate whether scDNAs could induce autophagosome formation, EGFP-LC3–expressing cells were transfected with HT-DNA or scDNAs (1 µg/mL). Six hours after transfection, cells were fixed and imaged. To investigate the effect of scDNAs on autophagic flux, cells were pre-treated with 20 μM chloroquine (MedChemExpress) 1 h before DNA transfection. To calculate the average number of EGFP-LC3–positive dots per cell, a minimum of 200 cells were counted in at least three independent experiments.

#### p40PX–EGFP puncta imaging

RAW264.7, HEK293T-cGAS, and HEK293T-Vector cells were transfected with p40PX–EGFP plasmid and stable clones expressing p40PX-EGFP fusion protein were selected with G418 (RAW264.7 cells: 200 µg/mL; HEK293T-cGAS and HEK293T-Vector cells: 1 mg/mL). p40phox PX-EGFP–expressing cells were transfected with HT-DNA or scDNAs (1 µg/mL). Six hours after transfection, cells were fixed and imaged. To calculate the average number of p40PX-EGFP puncta per cell, at least 200 cells were counted in at least three independent experiments.

#### Cytosolic dsDNA imaging

Cells were exposed to UV irradiation (200 J/m^2^), camptothecin (MedChemExpress), etoposide, or adriamycin (MedChemExpress), and cytosolic dsDNA was stained using anti-double-stranded DNA monoclonal antibody (listed in Extended Data Table 4) 4 h after UV irradiation or 24 h after exposure to camptothecin, etoposide, and adriamycin.

### Electron microscopy

RAW264.7 cells were transfected with scDNAs (1 µg/mL) or mock-transfected. Six hours after transfection, cells were fixed, sectioned, stained, and coated at the Electron Microscopy Core Facility of Chongqing Medical University (Chongqing, China). Images were visualized using a Hitachi JEM-1400 Plus electron microscope (Hitachi Koki Co., Ltd., Tokyo, Japan).

### Reverse transcription and quantitative PCR

Total RNA was prepared using Trizol reagent (Life Technologies), incubated with RNase-free DNase I (Promega, Madison, WI, USA) for 30 min, and reverse-transcribed using HiScript III-RT SuperMix for qPCR (Vazyme Biotech Co., Ltd.). SYBR Green real-time PCR was performed using the ChamQ Universal SYBR qPCR master mix (Vazyme Biotech Co., Ltd.) and ABI 7500 FAST sequence detection system (Life Technologies). Primer sequences are listed in Extended Data Table 1.

### Statistical analysis

All statistical analyses were performed using Student’s *t*-tests via SPSS software (v.22.0; IBM Corp., Armonk, NY, USA). All results are presented as the mean ± standard error of the mean of at least three independent experiments. In all experiments, the significance level was set as α = 0.05, and p < 0.05 indicates significant intergroup differences.

## Data availability statement

All data needed to evaluate the conclusions in the paper are included in the paper and/or the Supplementary Materials. The sequencing data of scDNAs are deposited in the NCBI Sequence Read Archive under BioSample accession: SAMN29261763, SAMN29261764, and SAMN29265103.

## Code availability

The code for sequencing analysis is available on GitHub at https://github.com/DrTangkaifu/scDNA-sequencing-analysis-script.

## Acknowledgments

We thank Dr. Peng Huang at the Electron Microscopy Core Facility of Chongqing Medical University for training in electron microscopy sample preparation and image processing. This work was supported by the National Natural Science Foundation of China (81972648, 82172915, 81773011) and CQMU Program for Youth Innovation in Future Medicine (W0084).

## Author contributions

K.F.T. conceived and designed the study, analyzed the data, and wrote the manuscript. Y.L., X.C. Y.Z., X.W., L.C., W.W., S.Z., M.H., Z.D., X.C., J.Z., H.L., D.M., X.W., H.J. and Y.W.L. performed the experiments. J.J. and S.H. performed the bioinformatic analysis. K.F.T. and A.H. supervised the study.

## Supplementary information

Supplementary Information is available for this paper.

## Corresponding author

Correspondence and requests for materials should be addressed to Kai-Fu Tang.

## Peer reviewer information

Reprints and permissions information is available at www.nature.com/reprints.

