## Extended Data Tables and figure legends for "Small cytosolic dsDNAs repress cGAS activation and induce autophagy"

Extended Data Table 1. The oligonucleotide sequences used in this study

| **Gene name** | | **sequence (5'-3')** |
|---|---|---|
| Human IFNβ primer | Forward | TCTCCTCAGGGATGTCAAAG |
| | Reverse | CAACAAGTGTCTCCTCCAAAT |
| Mouse IFNβ primer | Forward | TCCTGCTGTGCTTCTCCACCACA |
| | Reverse | AAGTCCGCCCTGTAGGTGAGGTT |
| Human GAPDH primer | Forward | ACCCACTCCTCCACCTTTG |
| | Reverse | CACCACCCTGTTGCTGTAG |
| Mouse GAPDH primer | Forward | ACGGCCAAATCCGTTCACACC |
| | Reverse | ACGGCCGCATCTTCTTGTGCA |
| Human p62 primer | Forward | GAGAAGCCCTCAGACAGGTG |
| | Reverse | GTGTCCGTGTTTCACCTTCC |
| Mouse p62 primer | Forward | TCCGATTCTGGCATCTGTAG |
| | Reverse | GCCAAAGTGTCCATGTTTCA |
| Teasy primer* | Forward | CGGCCAGTGAATTGTAATACG |
| | Reverse | CTCAAGCTATGCATCCAACG |
| P24 | Sense | GAAGTCAGGACGTGGCACGAAGAA |
| | Antisense | TTCTTCGTGCCACGTCCTGACTTC |

* The Teasy primer was used for qPCR to quantify the spike-in DNA, and for PCR amplification of the scDNAs ligated into the pGEM®-T Easy Vector.

Extended Data Table 2. The plasmids used in this study

| **name** | **Vendor** | **Catalog number** |
|---|---|---|
| pcDNA4-Beclin1(FL) | Addgene | #24388 |
| pCMV-Myc-BECN1 | Miaolingbio | #P6909 |
| pLenti-CMV-cGAS-HA | Addgene | #130910 |
| pcDNA4-Vps34-Flag | Addgene | #24398 |
| p40PX-EGFP | Addgene | #19010 |
| pMD2.G | Addgene | #12259 |
| psPAX2 | Addgene | #12260 |
| pGEX6P1-hcGAS（157-522） | Addgene | #108676 |
| pEGFP-N1-cGAS | Inovogen Tech. Co. | Custom-made |

Extended Data Table 3. The siRNA sequences used in this study.

| **Gene name** | | **sequence (5'-3')** |
|---|---|---|
| sicGAS-1 | Sense | GGAUUGAGCUACAAGAAUAdTdT |
| | Antisense | UAUUCUUGUAGCUCAAUCCdTdT |
| sicGAS-2 | Sense | GAUUGAGCUACAAGAAUAUdTdT |
| | Antisense | AUAUUCUUGUAGCUCAAUCdTdT |
| siCon | Sense | AAUUCUCCGAACGUGUCACGUdTdT |
| | Antisense | ACGUGACACGUUCGGAGAAUUdTdT |

Extended Data Table 4. The antibodies used in this study

| **Antibodies** | **Vendor** | **Catalog number** | **Application** | **Dilution** | **Secondary antibodies** |
|---|---|---|---|---|---|
| Flag | proteintech | 66008-2-Ig | WB | 1:1000 | Mouse |

| | | | | | |
|---|---|---|---|---|---|
| HA | GeneTex | GTX115044 | WB | 1:1000 | Rabbit |
| Rubicon | Abcam | ab92388 | WB | 1:1000 | Rabbit |
| Beclin1 | proteintech | 11306-1-AP | WB | 1:1000 | Rabbit |
| P62/SQSTM1 | proteintech | 18420-1-AP | WB | 1:1000 | Rabbit |
| Phospho-IRF3 (Ser396) | Cell Signaling Technology | #29047S | WB | 1:1000 | Rabbit |
| Mouse Phospho-STING (Ser365) | Cell Signaling Technology | #72971S | WB | 1:1000 | Rabbit |
| Human Phospho-STING (Ser366) | Cell Signaling Technology | # 50907S | WB | 1:1000 | Rabbit |
| LC3 | proteintech | 14600-1-AP | WB | 1:1000 | Rabbit |
| Mouse cGAS | Cell Signaling Technology | #31659 | WB | 1:1000 | Rabbit |
| Human cGAS | Cell Signaling Technology | #15102 | WB | 1:1000 | Rabbit |
| GAPDH | Cell Signaling Technology | #2118S | WB | 1:3000 | Rabbit |
| Anti-DNA Antibody, double stranded, clone AE-2 | Millipore | MAB1293 | IF | 1:100 | Mouse |
| Goat anti-rabbit IgG(H+L) secondary antibody,HRP conjugate | Biosharp | BL003A | WB | 1:10000 | |
| Goat anti-mouse IgG(H+L) secondary antibody,HRP conjugate | Biosharp | BL001A | WB | 1:10000 | |
| Goat Anti-mouse IgG (H+L) secondary antibody, FITC conjugated | BOSTER | BA1101 | IF | 1:50 | |

### Extended data figure legends

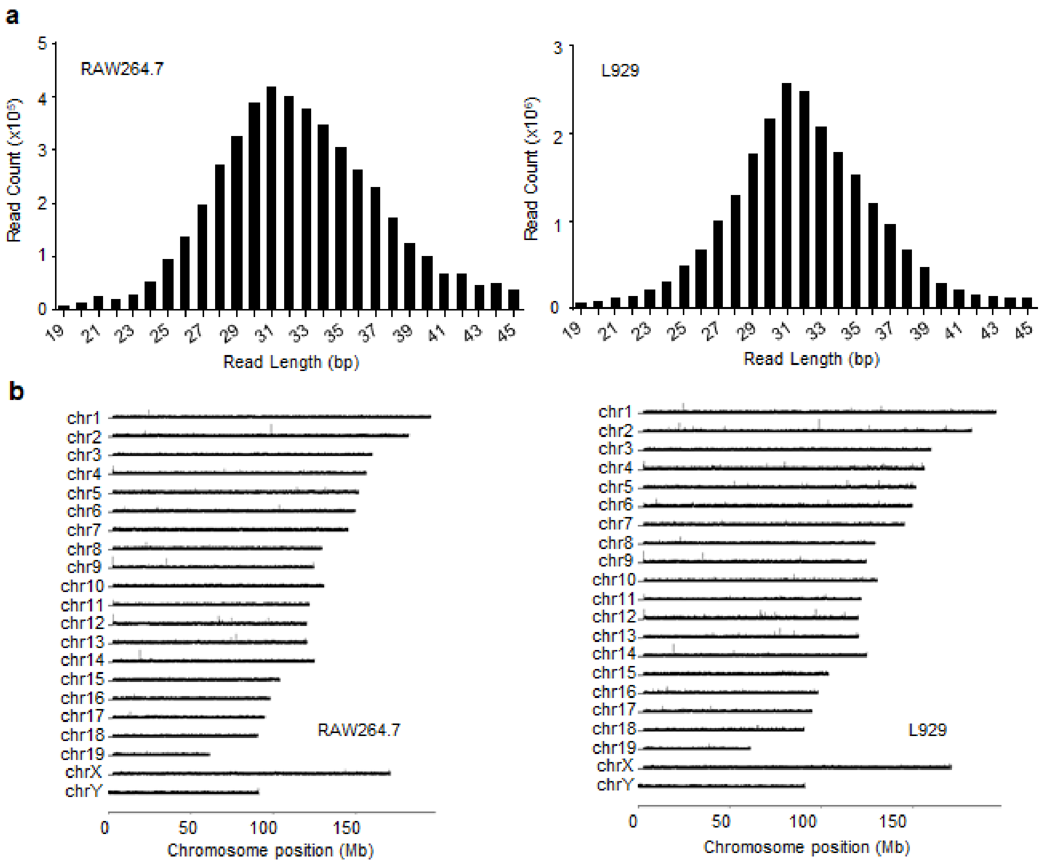


**Extended Data Figure 1. Sequencing of small cytosolic DNAs (scDNAs) purified from different cell lines.**

**a,** Read length distribution of the sequenced scDNAs purified from RAW264.7 and L929 cells. Only the sequences that could be mapped to the genome are shown. **b,** Density of sequenced reads along different chromosomes plotted as $\log_2$ (10 kilobases per million mapped reads).

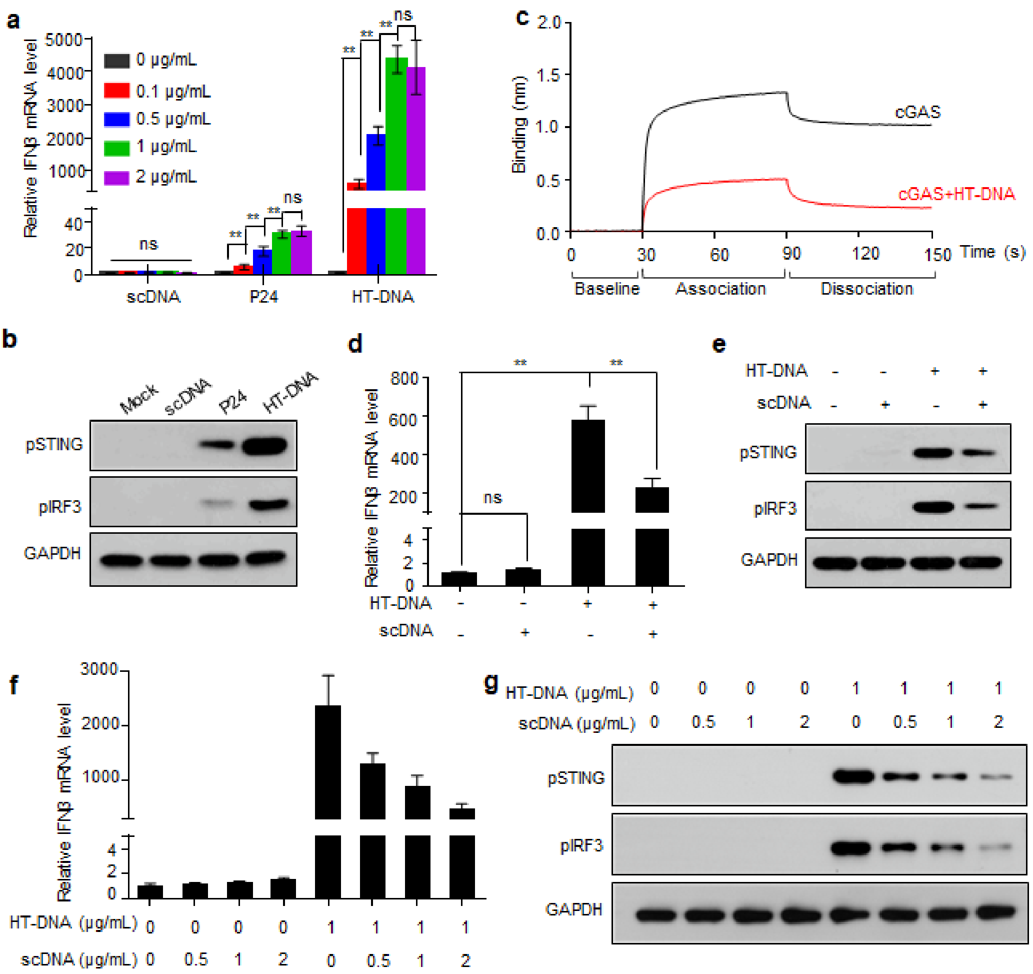


**Extended Data Figure 2. scDNAs compete with long dsDNA to bind cyclic GMP-AMP synthase (cGAS) and inhibit long dsDNA-induced cGAS activation.**

**a,** L929 cells were transfected with scDNAs, a 24 bp dsDNA (P24), or herring testis DNA (HT-DNA) at different concentrations, and interferon β (*IFNβ*) mRNA levels were quantified. **b,** L929 cells were transfected with scDNAs, P24, or HT-DNA (1 μg/mL) or mock-transfected, and phosphorylated (p) stimulator of interferon genes (STING) and p-interferon regulatory factor 3 (IRF3) levels were determined. **c,** Competition of scDNAs and HT-DNA for binding to cGAS. Biotin-labeled scDNAs were immobilized on a streptavidin-coated sensor and then loaded with cGAS alone (50 ng/μL) or cGAS pre-incubated with unlabeled HT-DNA (300 ng/μL). **d, e,** THP-1 cells were transfected with HT-DNA (1 μg/mL) or scDNAs (1 μg/mL) purified from L929 cells or co-transfected with both HT-DNA (1 μg/mL) and scDNAs (1 μg/mL), and *IFNβ* mRNA (d), pSTING, and pIRF3 (e) levels were determined. **f, g,** L929 cells were transfected with HT-DNA (1 μg/mL) or different doses of scDNAs purified from THP-1 cells or co-transfected with HT-DNA (1 μg/mL) and different doses of scDNAs, and *IFNβ* mRNA (f), pSTING, and pIRF3 (g) levels were determined. Data (a, d and f) are expressed as mean ± standard error of the mean (s.e.m) of three biological replicates. Statistical analysis was performed using two-sided Student's *t*-tests; ** indicates $p < 0.01$; ns indicates $p > 0.05$.

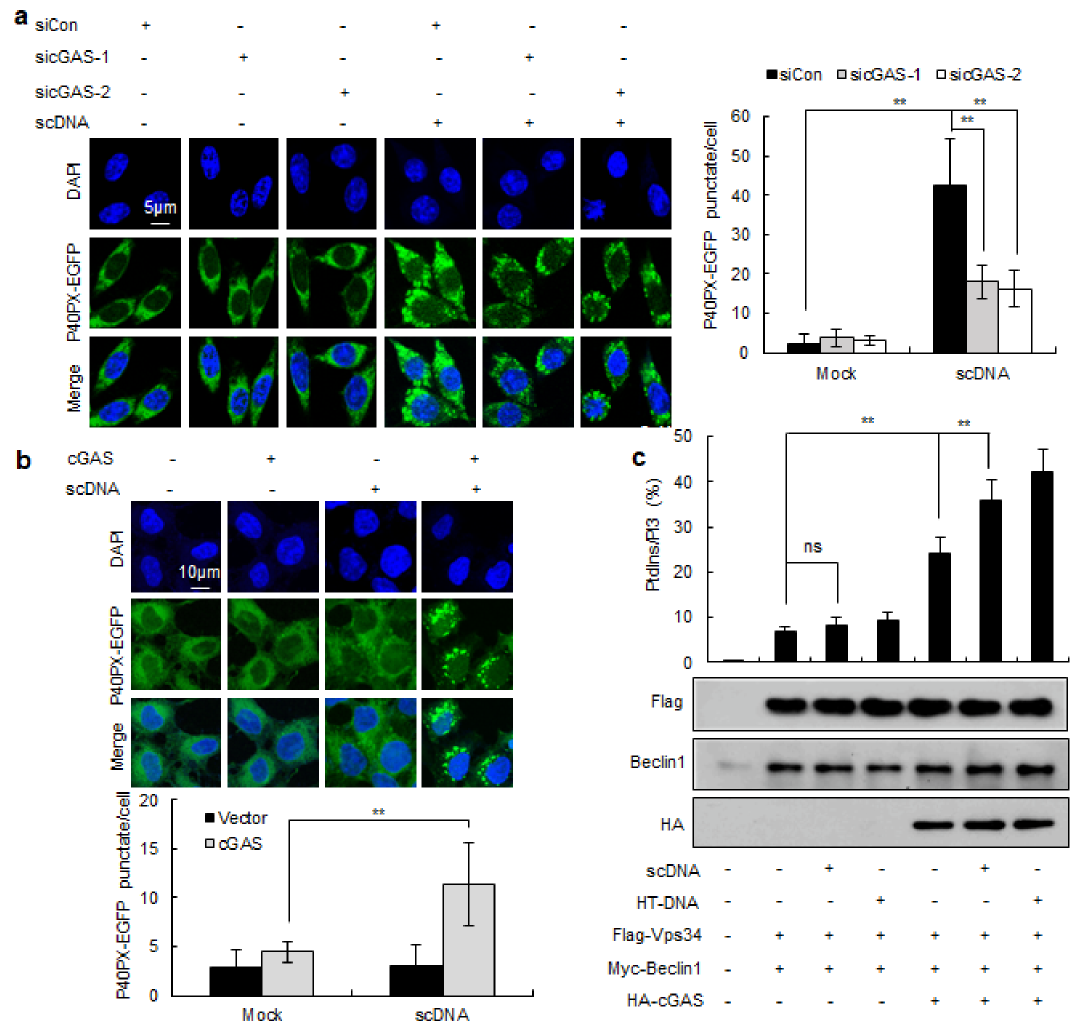


**Extended Data Figure 3. scDNAs promote phosphatidylinositol 3-kinase class III (PI3KC3) kinase activity in a cGAS-dependent manner.**

**a,** RAW264.7 cells stably expressing p40PX-EGFP were transfected with cGAS siRNAs or control siRNAs and stimulated with scDNAs (1 μg/mL) or mock-transfected. The number of p40PX-EGFP foci was determined using confocal microscopy. Data are expressed as mean ± s.e.m of three biological replicates, $n \geq 200$ cells. Statistical analysis was performed using two-sided Student's *t*-tests; ** indicates $p < 0.01$. **b,** HEK293T cells stably expressing p40PX-EGFP alone or together with cGAS were stimulated with scDNAs (1 μg/mL) or mock-transfected. The number of p40PX-EGFP foci was determined using confocal microscopy. Data are expressed as mean ± s.e.m of five biological replicates, $n \geq 200$ cells. Statistical analysis was performed using two-sided Student's *t*-tests; ** indicates $p < 0.01$. **c,** Flag-PI3KC3 and Myc-Beclin-1 plasmids were co-transfected into HEK293T-cGAS and HEK293T-Vector cells, and the cells were stimulated with scDNAs or HT-DNA (1 μg/mL). Flag-PI3KC3 complexes were immunoprecipitated using anti-Flag M2 affinity gel and subjected to an *in vitro* PI3KC3 kinase assay. HT-DNA was used as a positive control. Data are expressed as mean ± s.e.m of three biological replicates. Statistical analysis was performed using two-sided Student's *t*-tests; ** indicates $p < 0.01$; ns indicates $p > 0.05$.

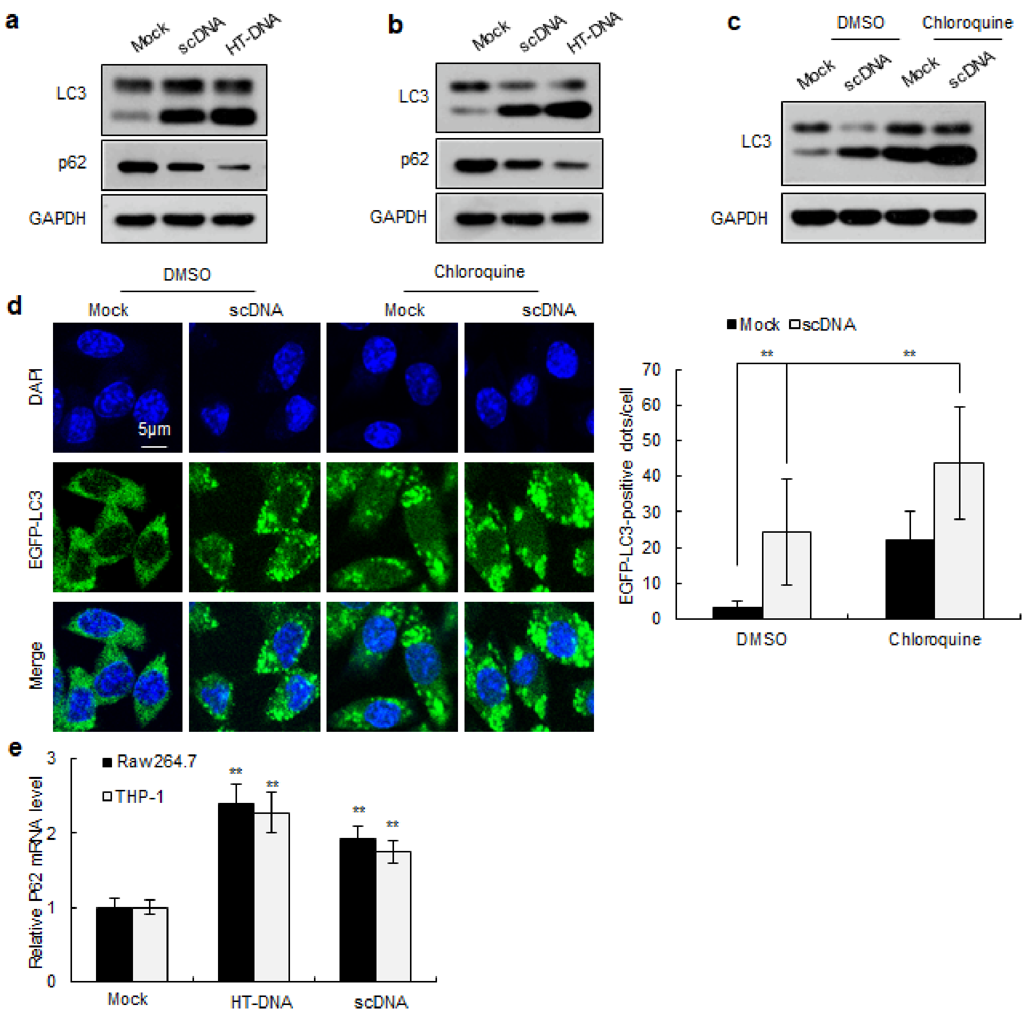


**Extended Data Figure 4. scDNAs induce autophagy in human and mouse cell lines.**

**a,** RAW264.7 cells were transfected with scDNAs purified from THP-1 cells (1 μg/mL) or mock-transfected, and LC3-II and p62 levels were determined. **b,** THP-1 cells were transfected with scDNAs purified from RAW264.7 cells (1 μg/mL) or mock-transfected, and LC3-II and p62 levels were determined. HT-DNA was used as a positive control in a and b. **c,** RAW264.7 cells were transfected with scDNAs (1 μg/mL) in the presence or absence of chloroquine, and LC3-II levels were determined. **d,** RAW264.7 cells stably expressing EGFP-LC3 were transfected with scDNAs (1 μg/mL) or mock-transfected in the presence or absence of chloroquine, and the formation of LC3-EGFP puncta was analyzed using confocal microscopy. Data are expressed as mean ± s.e.m of three biological replicates, $n \geq 200$ cells. Statistical analysis was performed using two-sided Student's *t*-tests; ** indicates p < 0.01. **e,** Quantification of *p62* mRNA levels in different cell lines transfected with scDNAs or HT-DNA (1 μg/mL). Data are expressed as mean ± s.e.m of three biological replicates. Statistical analysis is performed using two-sided Student's *t*-tests; ** indicates p < 0.01.

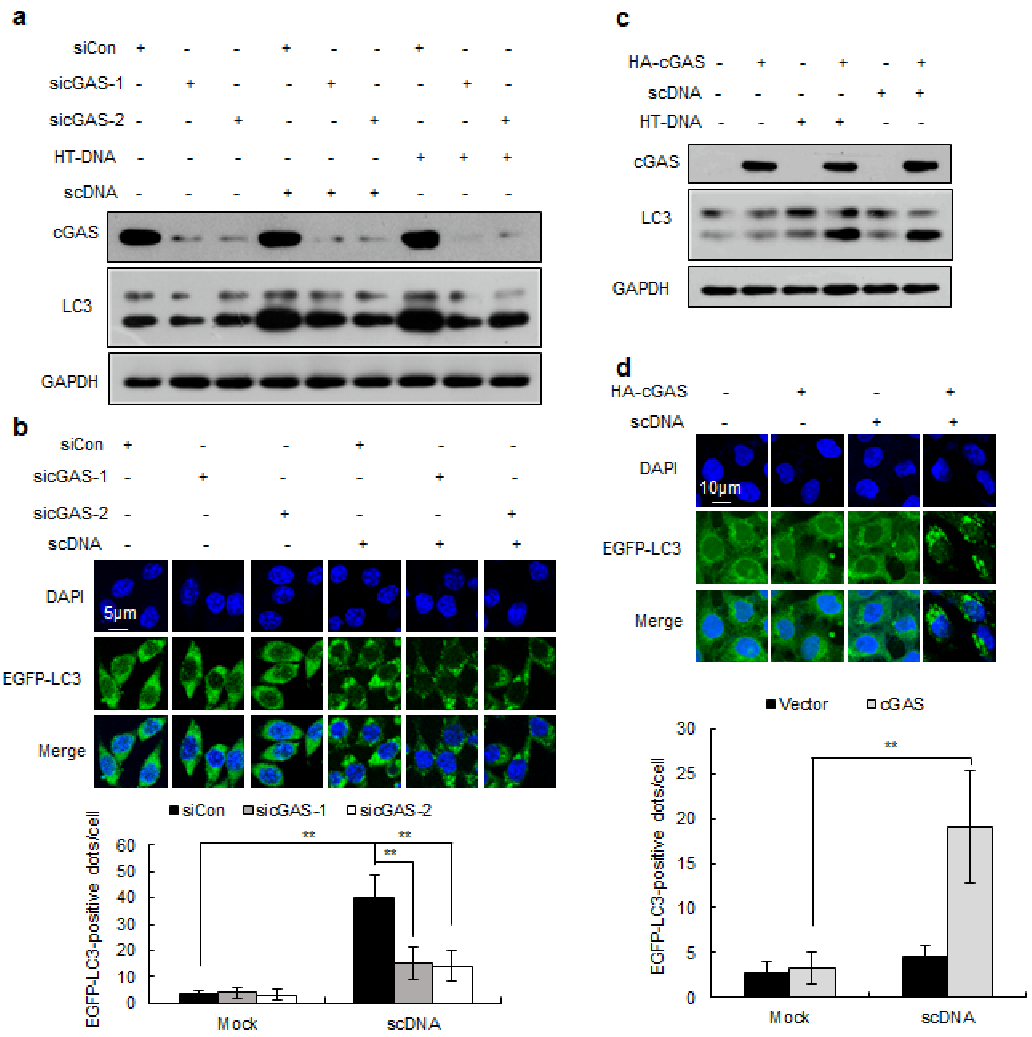


**Extended Data Figure 5. cGAS is required for scDNA-induced autophagy.**

**a,** RAW264.7 cells were transfected with control or cGAS siRNAs and stimulated with or without scDNAs (1 μg/mL). Whole cell lysates were harvested and analyzed by immunoblotting using the indicated antibodies. HT-DNA was used as a positive control. **b,** RAW264.7 cells stably expressing EGFP-LC3 were transfected with control or cGAS siRNAs and then transfected with scDNAs (1 μg/mL) or mock-transfected and analyzed using confocal microscopy. Data are expressed as mean ± s.e.m of four biological replicates, $n \geq 200$ cells. Statistical analysis was performed using two-sided Student's *t*-tests; ** indicates $p < 0.01$. **c,** HEK293T-Vector or HEK293T-cGAS cells were stimulated with or without scDNAs (1 μg/mL). Whole cell lysates were harvested and analyzed by immunoblotting using the indicated antibodies. HT-DNA was used as a positive control. **d,** HEK293T cells expressing EGFP-LC3 alone or in combination with cGAS were stimulated with scDNAs (1 μg/mL) or mock-transfected and analyzed using confocal microscopy. Data are expressed as mean ± s.e.m of three biological replicates, $n \geq 200$ cells. Statistical analysis was performed using two-sided Student's *t*-tests; ** indicates $p < 0.01$.

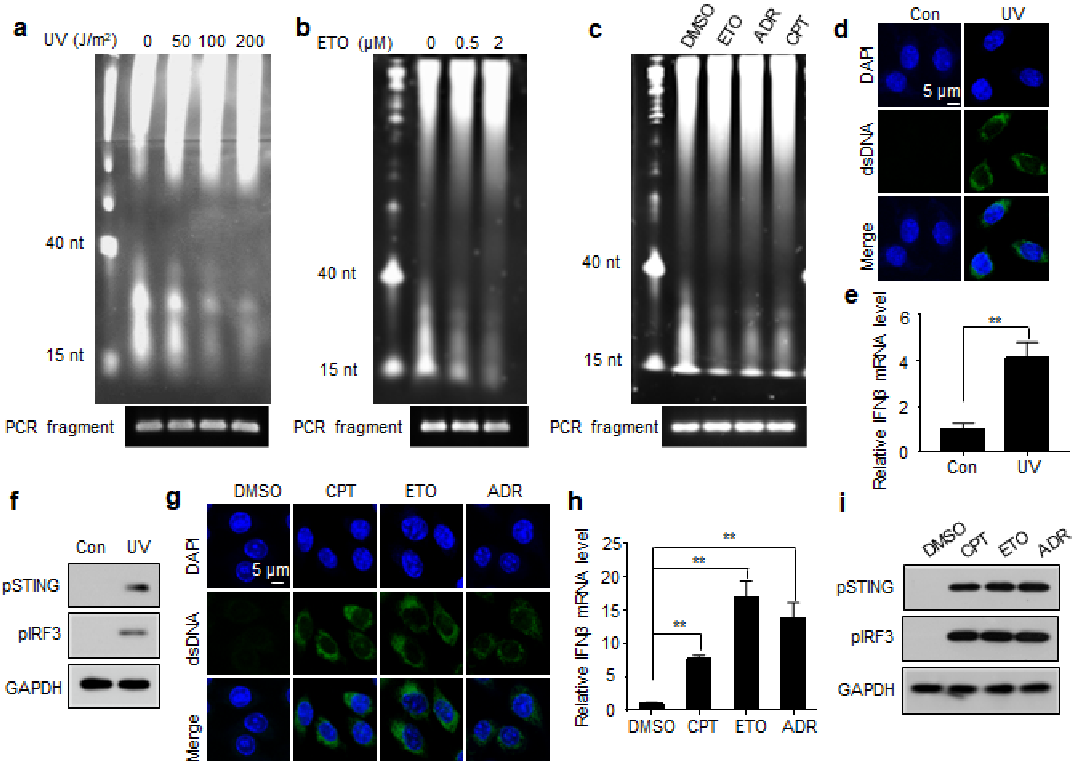


**Extended Data Figure 6. DNA-damaging agents reduce scDNA levels, induce the presence of cytosolic dsDNA, and lead to cGAS-STING activation.**

**a,** RAW264.7 cells were exposed to different doses of ultraviolet (UV) irradiation, and cytosolic DNA was purified and subjected to PAGE analysis. **b,** RAW264.7 cells were treated with different doses of etoposide (ETO), and cytosolic DNA was subjected to PAGE analysis. **c,** L929 cells were treated with etoposide (2 µM), adriamycin (ADR, 5 µM), or camptothecin (CPT, 0.2 µM), and cytosolic DNA was subjected to PAGE analysis. **d–f,** RAW264.7 cells were exposed to 200 J/m$^2$ UV irradiation, and cytosolic dsDNA (d), *IFNβ* mRNA (e), pSTING, and pIRF3 (f) levels were determined. **g–i,** RAW264.7 cells were treated with camptothecin (0.2 µM), etoposide (2 µM), or adriamycin (5 µM), and cytosolic dsDNA (g), *IFNβ* mRNA (h), pSTING and pIRF3 (i) levels were determined. The PCR products of spike-in DNA were used as loading controls (a–c). Data (e and h) are expressed as mean ± s.e.m of three biological replicates. Statistical analysis was performed using two-sided Student's *t*-tests; ** indicates $p < 0.01$.

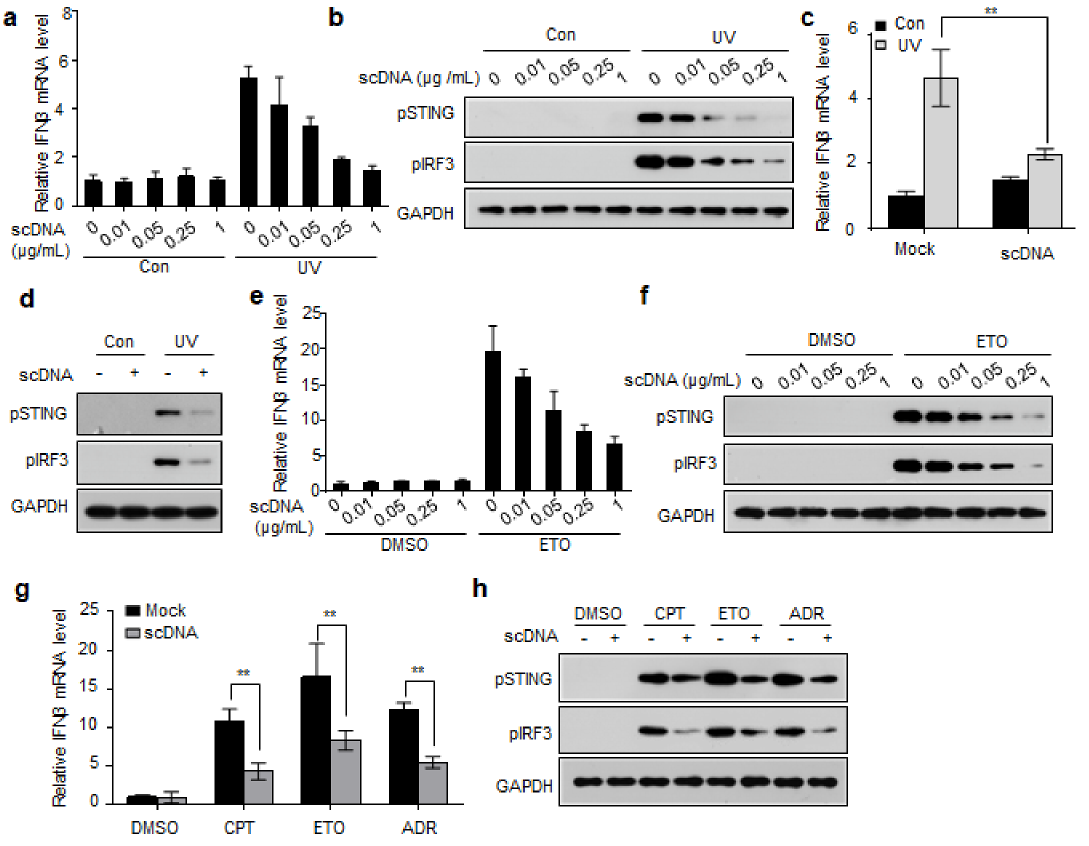


**Extended Data Figure 7. scDNA transfection partially blocks cGAS-STING activation following DNA damage.**

**a, b,** RAW264.7 cells were transfected with different doses of scDNAs or mock-transfected and exposed to 200 J/m$^2$ UV irradiation. *IFNβ* mRNA (a), pSTING, and pIRF3 (b) levels were determined. **c, d,** L929 cells were transfected with scDNAs (0.25 μg/mL) or mock-transfected and exposed to 200 J/m$^2$ UV irradiation, and *IFNβ* mRNA (c), pSTING, and pIRF3 (d) levels were determined. **e, f,** RAW264.7 cells were transfected with different doses of scDNAs or mock-transfected and exposed to etoposide (2 μM). *IFNβ* mRNA (e), pSTING, and pIRF3 (f) levels were determined. **g, h,** L929 cells were transfected with scDNAs (0.25 μg/mL) or mock-transfected and exposed to camptothecin (0.2 μM), etoposide (2 μM), or adriamycin (5 μM). *IFNβ* mRNA (g), pSTING, and pIRF3 (h) levels were determined. Data (a, c, e and g) are expressed as mean ± s.e.m of three biological replicates. Statistical analysis was performed using two-sided Student's *t*-tests; ** indicates $p < 0.01$.

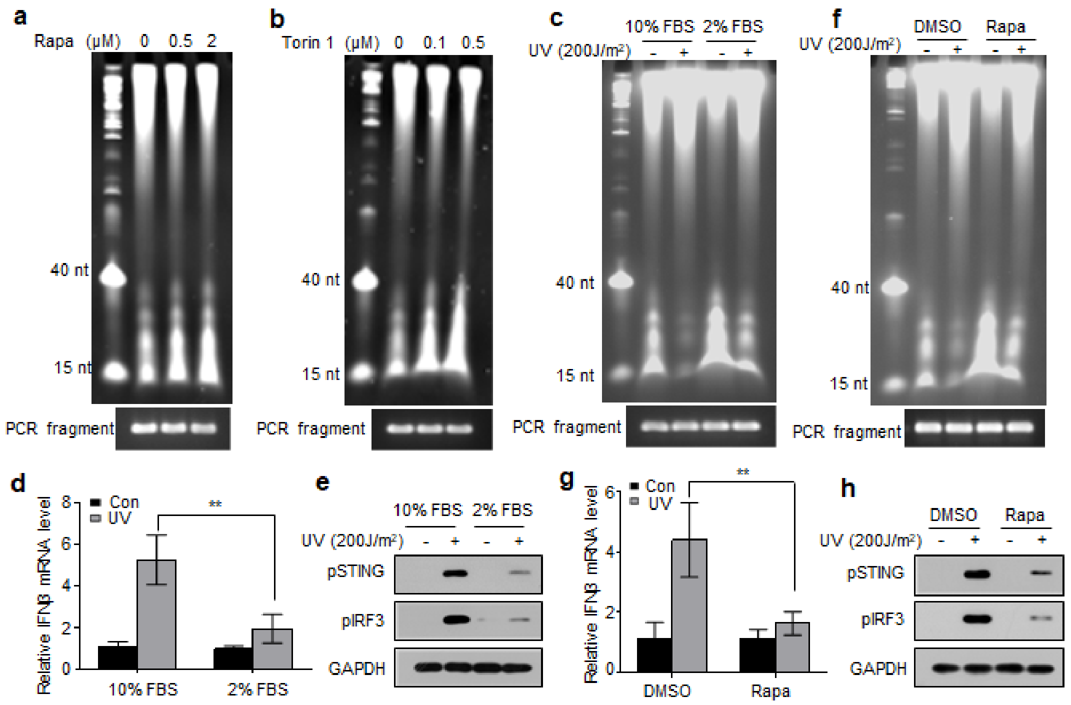


**Extended Data Figure 8. Treatment with autophagy inducers increases scDNA levels and blocks cGAS-STING activation following DNA damage.**

**a,** L929 cells were treated with different doses of rapamycin, and cytosolic DNA was purified and subjected to PAGE analysis. **b,** L929 cells were treated with different doses of Torin 1, and cytosolic DNA was purified and subjected to PAGE analysis. **c,** RAW264.7 cells were exposed to 10% or 2% FBS and irradiated with 200 J/$m^2$ UV or unirradiated, and cytosolic DNA was purified and subjected to PAGE analysis. **d, e,** RAW264.7 cells were exposed to 10% or 2% FBS and irradiated with 200 J/$m^2$ UV or unirradiated, and *IFNβ* mRNA (d), pSTING, and pIRF3 (e) levels were determined. **f,** RAW264.7 cells were treated with rapamycin (2 μM) and irradiated with 200 J/$m^2$ UV or unirradiated, and cytosolic DNA was purified and subjected to PAGE analysis. **g, h,** RAW264.7 cells were treated with rapamycin (2 μM) and irradiated with 200 J/$m^2$ UV or unirradiated, and *IFNβ* mRNA (g), pSTING, and pIRF3 (h) levels were determined.

The PCR products of spike-in DNA were used as loading controls (a–c, f). Data (d and g) are expressed as mean ± s.e.m of three biological replicates. Statistical analysis was performed using two-sided Student's *t*-tests; ** indicates $p < 0.01$.
